## Supplementary Figures for "Variable Induction of Pro-inflammatory Cytokines by Commercial SARS CoV-2 Spike Protein Reagents: Potential Impacts of LPS on *In Vitro* Modeling and Pathogenic Mechanisms *In Vivo*": Ouyang W., et al. SARS CoV-2 Spike Protein and LPS - Supplementary Information.pdf

**Supplementary Figure 1**

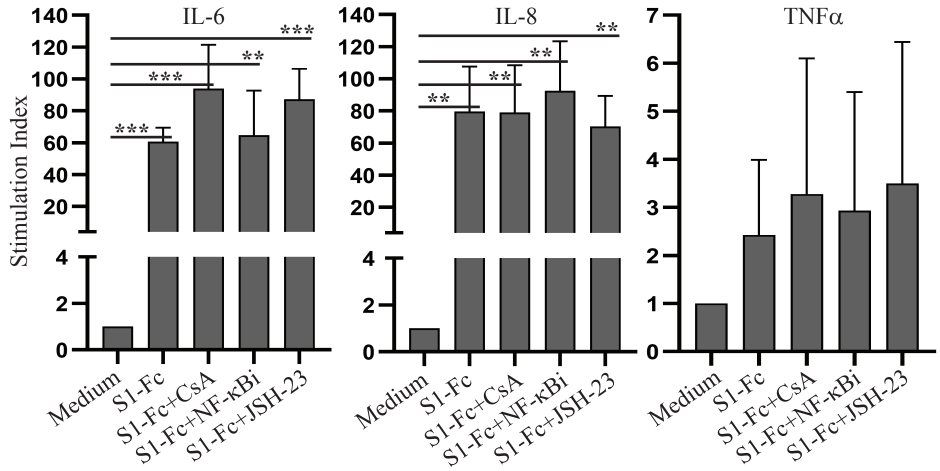


Blockade of the NFAT and NF-κB signaling pathways does not inhibit S1-Fc-induced cytokine production. Rested PBMC were cultured with or without 2.0 µg/mL of S1-Fc from Vendor#2 in the presence or absence of cyclosporin A (CSA), NF-κB SN50 cell permeable inhibitory peptide (NF-κBi) or NF-κB inhibitor JSH-23 for 24 hours. The levels of IL-6 and TNFα were measured using the CBA human inflammatory cytokine kit and flow cytometric analysis. IL-8 concentrations were measured using an ELISA kit. The concentrations of the cytokines were calculated based on the standard curves. Data shown are statistical results of the stimulation indexes (mean ± SE) derived from 3 healthy donors.

**Supplementary Figure 2**

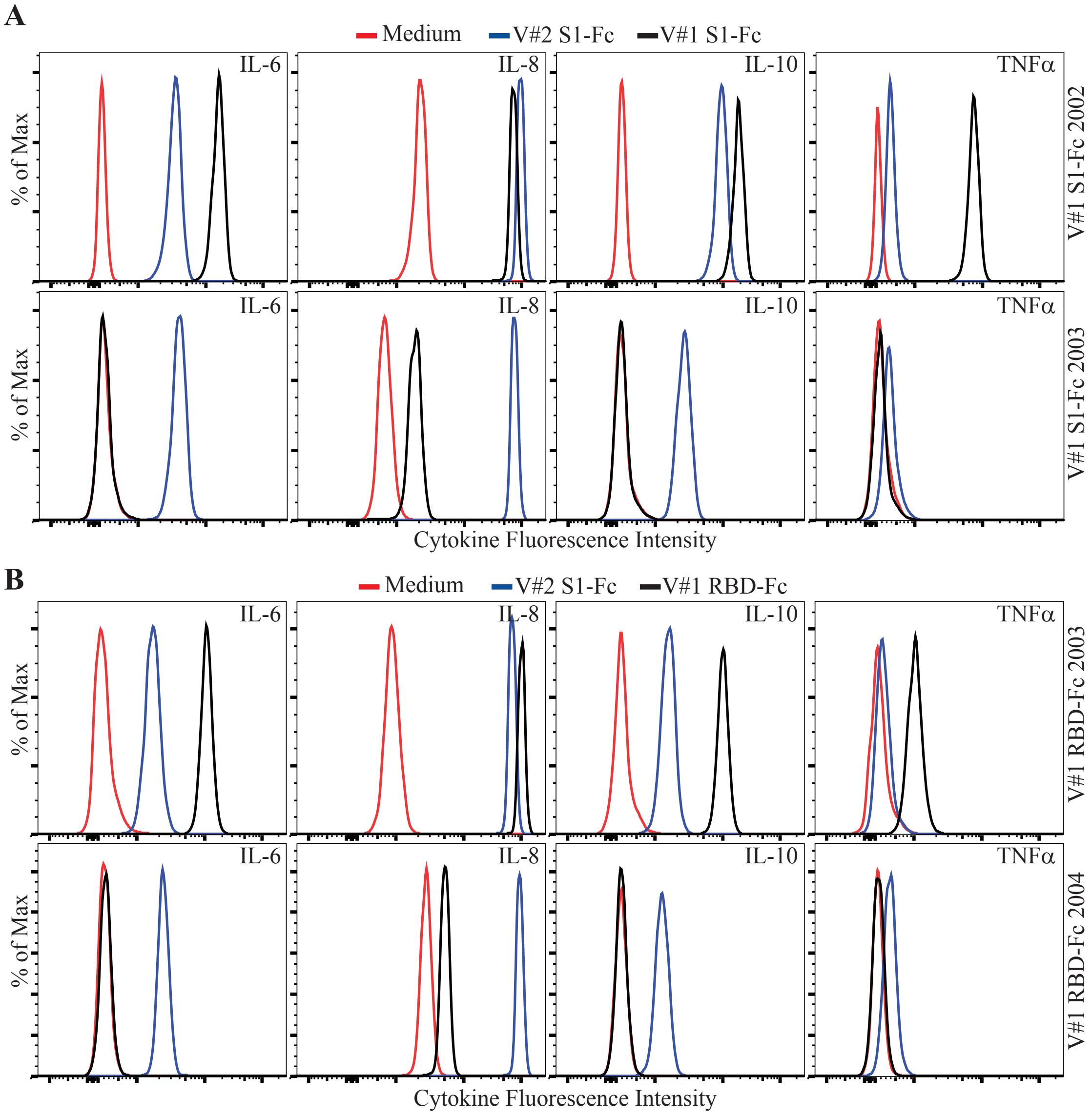


Batch differences in cytokine induction by lots of S1-Fc and RBD-Fc fusion proteins. Rested PBMC were cultured with or without 2.0 µg/mL of S1-Fc from Vendor #2 (V#2 S1-Fc), S1-Fc from Vendor #1 (lot 24056-2002-2 (V#1 S1-Fc 2002) and lot 24529-2003 (V#1 S1-Fc 2003)) (**A**), and RBD-Fc from Vendor #1 (lot 24530-2003 (V#1 RBD-Fc 2003) or lot 25130-2004 (V#1 RBD-Fc 2004)) (**B**) for 24 hours. The levels of IL-6, IL-8, IL-10 and TNFα were assessed using the CBA human inflammatory cytokine kit and flow cytometric analysis.

**Supplementary Figure 3**

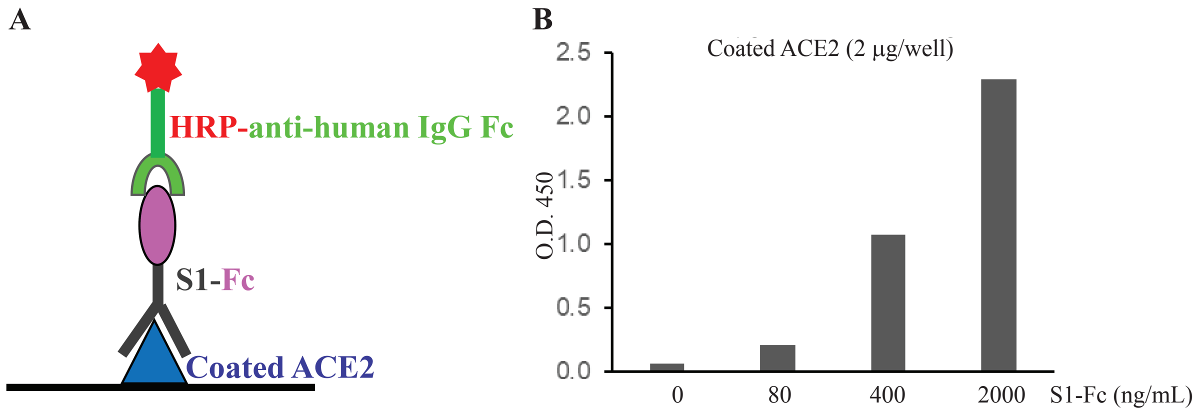


Development of an ELISA to measure the binding activity of spike protein to ACE2. (**A**) A schematic diagram showing the ELISA. (**B**) The binding activities of 5-fold serially diluted S1-Fc (from 2000 to 80 ng/mL) to plate-bound ACE2. The data are presented as absorbance at 450 nm (O.D. 450).

**Supplementary Figure 4**

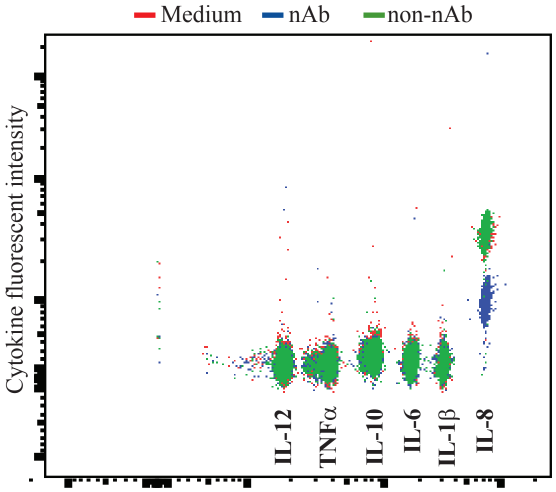


Anti-S1 antibodies do not induce cytokine production. Rested PBMC were cultured with or without neutralizing (nAb) or non-neutralizing (non-nAb) anti-S1 antibodies for 24 hours. The levels of IL-1β, IL-6, IL-8, IL-10, IL-12 and TNFα were assessed using the CBA human inflammatory cytokine kit and flow cytometric analysis.

**Supplementary Figure 5**

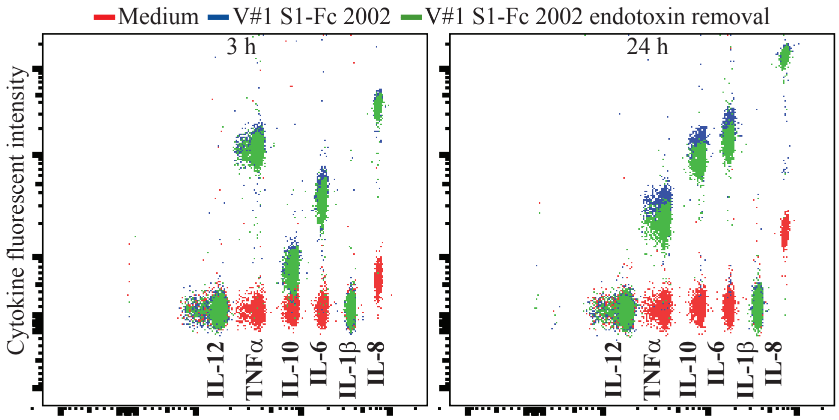


An endotoxin removal procedure fails to block pro-inflammatory potential of contaminated S1 fusion protein samples. Rested PBMC were cultured with or without S1-Fc (V#1 S1-Fc 2002) that was treated with or without endotoxin removal for 3 and 24 hours. The levels of IL-1β, IL-6, IL-8, IL-10, IL-12 and TNFα were assessed using the CBA human inflammatory cytokine kit and flow cytometric analysis.

**Supplementary Figure 6**

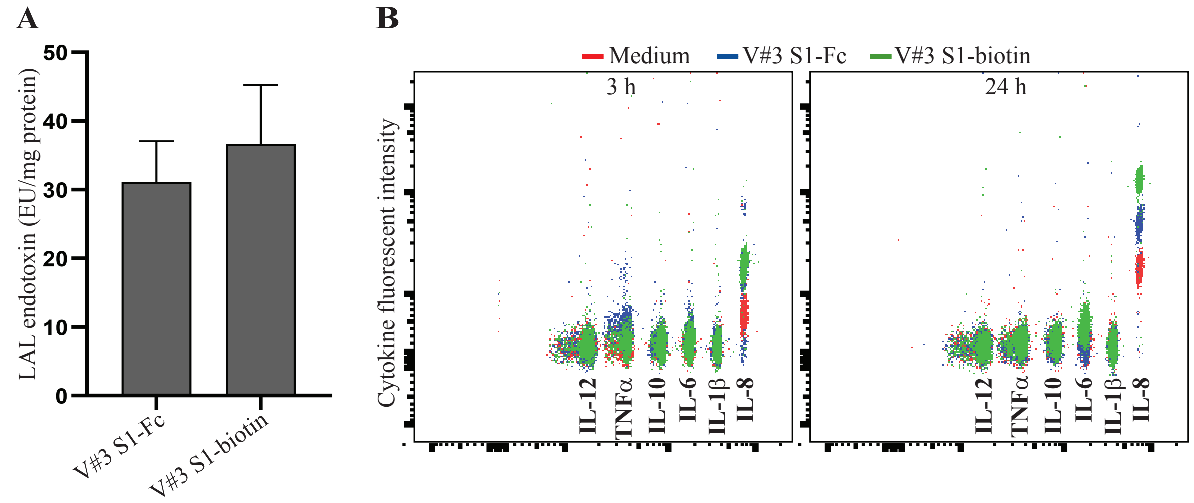


Spike protein preparations from Vendor#3 contain low levels of endotoxin and do not induce cytokine expression in human PBMC. (**A**) The levels of endotoxin in S1-Fc (V#3 S1-Fc) and S1-biotin (V#3 S1-biotin) from Vendor#3 were measured using the LAL Chromogenic Endotoxin Quantitation Kit. The concentrations of endotoxin were presented as EU/mg protein. Data shown are mean ± SE of the results from three independent experiments. (**B**) Rested PBMC were untreated or cultured with S1-Fc or S1-biotin from Vendor#3 for 3 and 24 hours. The levels of IL-1β, IL-6, IL-8, IL-10, IL-12 and TNFα were assessed using the CBA human inflammatory cytokine kit and flow cytometric analysis.

**Supplementary Figure 7**

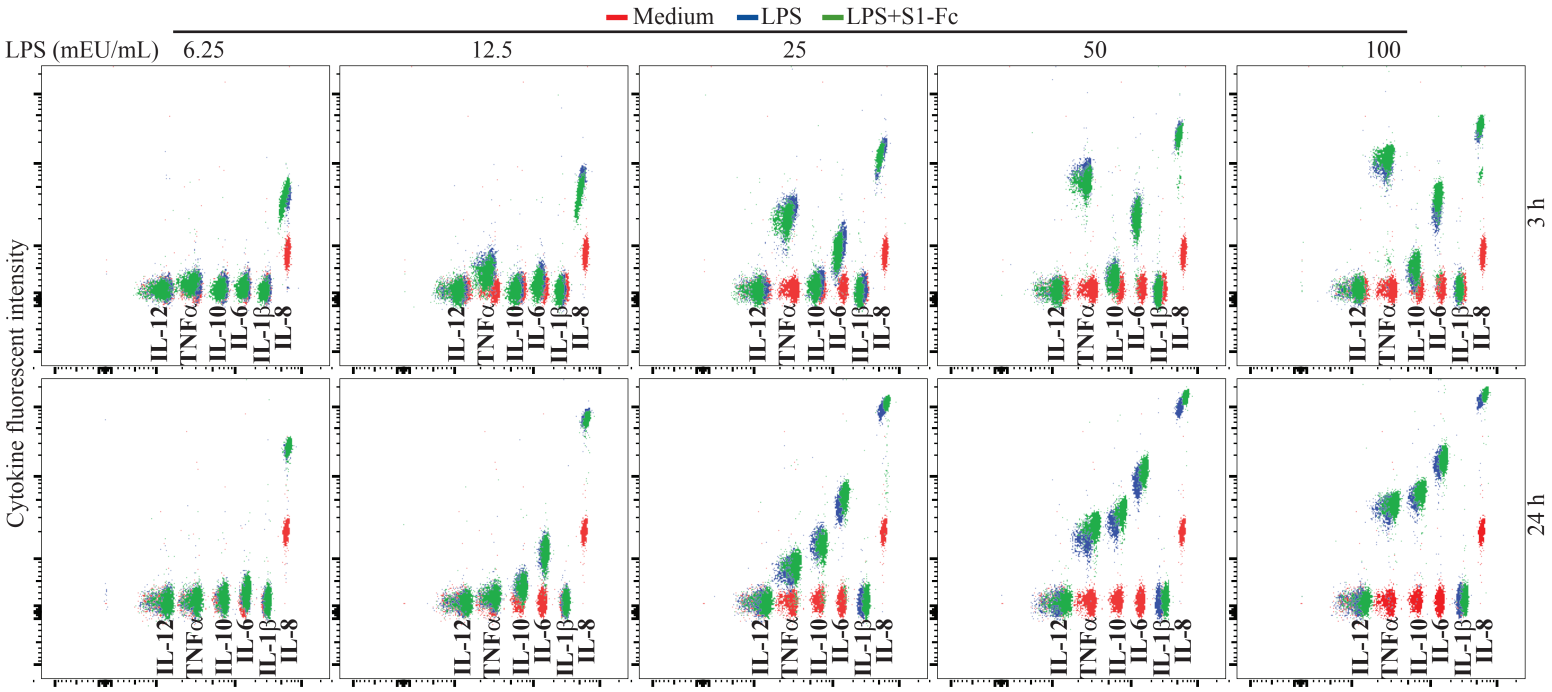


S1-Fc does not enhance LPS-induced cytokine production in human PBMC. Rested PBMC were stimulated with 6.25 - 100 mEU/mL of LPS in the presence or absence of 2 μg/mL of S1-Fc (Vendor#1, lot 24529-2003) for 3 and 24 hours. The levels of IL-1β, IL-6, IL-8, IL-10, IL-12 and TNFα were assessed using the CBA human inflammatory cytokine kit and flow cytometric analysis.

**Supplementary Figure 8**

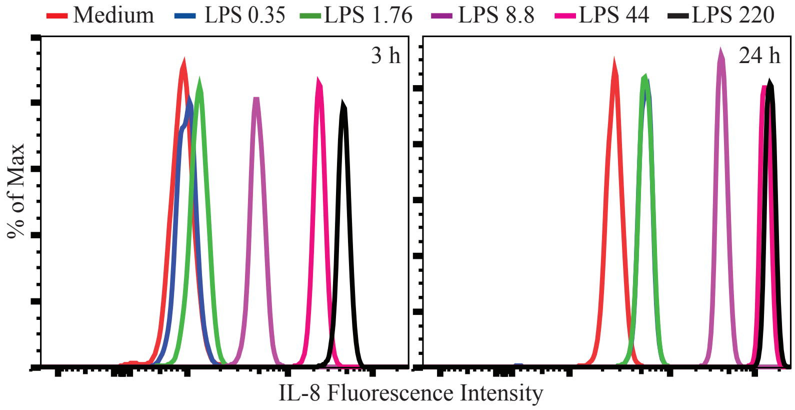


Dose-dependent IL-8 induction by LPS. Rested PBMC were untreated or cultured with 5-fold serially diluted LPS from 220 to 0.35 mEU/mL for 3 and 24 hours. The levels of IL-8 were assessed using the CBA human inflammatory cytokine kit and flow cytometric analysis.
